# Viral infection overcomes ineffectiveness of anti-tumoral CD8+ T cell mediated cytotoxicity

**DOI:** 10.1101/591198

**Authors:** Fan Zhou, Justa Kardash, Hilal Bhat, Vikas Duhan, Sarah-Kim Friedrich, Judith Bezgovsek, Halime Kalkavan, Michael Bergerhausen, Max Schiller, Yara Machlah, Tim Brandenburg, Cornelia Hardt, Michal Krolik, Lukas Flatz, Philipp A. Lang, Karl S. Lang

## Abstract

With the integration of PD-1 and CTLA-4 targeting immune checkpoint blockade into cancer treatment regimes, the anti-tumoral cytotoxicity of tumor-specific CD8^+^ T cells is well established. However, while the unresponsiveness of CD8^+^ T cells against big tumors is mainly explained by T cell exhaustion, other factors contributing to CD8^+^ T cell failure remain not well studied. Here we used a mouse melanoma model to study the interaction of growing tumor cells, innate immunity and CD8^+^ T cell responses induced by viral replication. Mouse model of melanoma (B16F10-OVA) and infections with arenaviruses. Growing B16F10-OVA cells did not induce systemic ablation of tumor specific CD8^+^ T cells. However, despite the presence of tumor-infiltrating CD8^+^ T cells, the anti-tumoral immune response was very limited. T cell anergy against the tumor was accompanied with a strong down-regulation of MHC-I on advanced tumors. LCMV infection restored the MHC class I expression, enhanced T cell function and lead to tumor regression. This study shows that tumor progression does not necessary lead to systemic exhaustion of the anti-tumoral CD8^+^ T cell response. Lack of innate signals is an additional reason for limited CD8^+^ T cell mediated cytotoxicity against the tumor.

## Introduction

Cancer development is strongly linked to accumulation of mutations that cause expression of high amount of tumor-specific and tumor-associated antigens. Such antigens could lead to the activation of an anti-tumoral immune responses, which might mediate control of the tumor growth[1-5]. Advanced neoplasms acquire several properties to escape from the host’s immune system leading to invasion of the tumor. Immune evasion mechanisms include dysregulation of the antigen presentation pathways, exhaustion of T cells, release of negative regulatory signals and recruitment of immunosuppressive cells to the tumor microenvironment supporting the establishment of the immune tolerance to cancer cells[6-11]. The immunogenic tolerance to tumors limits the effects of the anticancer chemotherapy and irradiation[12-14]. Therefore new approaches in anti-tumor therapy are needed to overcome the tumor immune escape.

Many human cancers contain infiltrating immune cells. In particular cytotoxic CD8^+^ T cells belong to the tumor microenvironment and play a key role in tumor eradication[4, 15-17]. Antigen-presenting cells (APCs) provide tumor-associated antigens to the cytotoxic T cells, leading to priming of such T cells[18, 19]. In fact, previous studies demonstrated that APCs, which are located in the lymph node (LN) internalize and cross-present tumor-associated antigens to cytotoxic CD8^+^ T cells[1, 20-22]. Beside the APCs, malignant cells are also able to trigger activation of cytotoxic T cells[23]. Presentation of tumor-specific neoantigens as well as tumor-associated antigens occur by MHC class I molecules expressed on the surface of tumor cells enabling cytotoxic T cells to recognize the antigen[23, 24]. In response to the detection of foreign antigens, cytotoxic T cells attack and destroy the MHC-I positive malignant cells. However, tumors use different escape mechanism to bypass the immunosurveillance by cytotoxic T cells. Loss or downregulation of MHC-I is a frequent mechanism allowing the neoplasms to evade identification by cytotoxic T cells[4, 23]. MHC-I downregulation was observed in a variety of malignancies revealing an association with progression or recurrence of disease, overall survival and increase in metastasis[25-27]. In particular, in tumors derived from epithelia, MHC-I molecules were reduced in more than 75% of patients[28]. Decrease of the MHC-I expression in tumors is a result of structural alterations in genes encoding MHC-I heavy chain, β-microglobulin or components of the antigen processing machinery[23, 29].

Viruses are well known to stimulate the innate and adaptive immune system. Recently, viral therapy has emerged as a promising approach to fight cancer. Cancer cells are a good target for non-oncolytic viruses which replicate selectively and activate immune defenses against tumor cells without harming normal tissue[30, 31]. In mouse models, previous research showed that acute infections with WE strain of lymphocytic choriomeningitis virus (LCMV-WE) exert antitumor effects[9, 32, 33]. Beside these principal inflammatory stimuli, LCMV can be used as a vaccine vector to immunize against tumor antigens[34-37].

The current research aimed to investigate the effect of viral infection with LCMV and Candid#1 on tumor-specific CD8^+^ T cells using a murine melanoma model.

## Materials and Methods

### Mice

C57BL6J (CD45.2^+^) and OT-1 were maintained on the C57BL/6 background. (back crossed as many as 16 times) and were bred as homozygotes. For adoptive transfer experiments mice, congenic for C57BL/6 (CD45.1) were used to distinguish between transferred cells (CD45.2) and endogenous (CD45.1) cells. The spontaneous tumor models B16F10-OVA (Melanoma) were used on C57BL/6 background. All animals were housed in single ventilated cages. Six- to eight-week-old, age and sex matched mice were used for all the experiments performed. All animals were housed in single ventilated cages. Animal experiments were authorized by the Nordrhein Westfalen Landesamt für Natur, Umwelt und Verbraucherschutz (Recklinghausen, Germany) and in accordance with the German law for animal protection or according to institutional guidelines at the Ontario Cancer Institute of the University Health Network and at McGill University.

### Viruses

The LCMV strain WE was originally obtained from F. Lehmann-Grube (Heinrich Pette Institute, Hamburg, Germany) and was propagated on L929 cells, MC57 cells, or both. LCMV-OVA was from Lukas Flatz (Department of Dermatology/Allergology, St. Gallen, Switzerland). Candid#1 was grown in Vero E6 cells derived from P. Cannon (Department of Molecular Microbiology and Immunology, University of Southern California, Los Angeles).

### Tumor induction

The C57BL/6-drived melanoma cell line B16F10-OVA-transfected clone was maintained at 37°C with 5% CO2 in DMEM medium supplemented with 10% heat-in-activated fetal calf serum, penicillin, streptomycin and Glutamine. 1×10^6^ B16F10-OVA cells were injected subcutaneously in 200ul Medium on the left flank. Tumor size was determined by the formula L×W×W where L=length, W=width, on the indicated days.

### In vivo killer assay

24 hours after B16-OVA injected subcutaneously to LCMV-OVA immunized mice, splenocytes differentially labeled with 20 or 200nM CFSE and load with or without 100nM ovalbumin_257-264_ peptide, respectively. The cells were collected 24 hr after i.v and s.c. injection. Splenocytes, which were not labeled with peptide served as negative control for determine the specific lysis.

### Flow cytometry

Tetramers were provided by the National Institutes of Health (NIH) Tetramer Facility. Cells were stained with allophycocyanin (APC)-labeled ovalbumin-(SIINFEKL) MHC class I tetramer (TCMetrix) for 15 minutes at 37°C. After incubation, the samples were stained with anti-CD8 (53-6.7) (eBioscience) for 30 minutes at 4°C. Absolute numbers of ova-specific CD8+ T cells were calculated from FACS analysis using fluorescing beads (BD Biosciences). All stained cells were analysed on an LSRII or a FACSFortessa (BD Biosciences) flow cytometer, and data were analysed with Flowjo software.

### Intracellular cytokine staining (ICS)

For ICS, Cells were stimulated with 100ng/ml PMA (Sigma) and 50 ng/ml ionomycin (Sigma-Aldrich) at 37°C for 3hr in the presence of Brefeldin A (10μg/ml; Sigma) to allow accumulation of intracellular cytokines. After staining of surface maker CD8, cells were fixed and permeabilized, followed by stain of IFN-γ (XMG 1.1).

### Histological analysis and Immunocytochemistry

Histological analysis were performed on snap-frozen tissue as previously described[30]. In brief, sections were fixed with acetone for 10 min, and non-specific antigens were blocked in phosphate-buffered saline (PBS) containing 2% FCS for 15 min, followed by various stainings for 45 min.

Immunocytochemistry was performed on cells grown on cover-slips[31]. In brief, cells were fixed in 4% Formalin/PBS for 10 min, washed in PBS and then permeabilized in 1% TritonX-PBS for 10 min. After blocking with 10% FCS/PBS for 30-60 min, various stainings for 45 min were followed. Cover slips were mounted on microscope slides using mounting medium (S3023, Dako).

Images were acquired with a fluorescence microscope (KEYENCE BZ II analyzer). Quantifications were performed using the Image J software (NIH).

### Statistical analysis

Graphs were compiled and statistical analyses were performed with Graphpad Prism software. Data are expressed as means ± SEM. Student’s t-test was used to detect statistically significant differences between groups. Significant differences between several groups were detected by one-way analysis of variance (ANOVA) with Bonferroni or Dunnett post hoc tests. The level of statistical significance was set at P < 0.05.

## Results

### Systemic expansion of tumor-specific CD8+ T cells despite growing tumors

To determine the influence of locally growing tumors on antigen specific CD8^+^ T cells, we treated C57BL/6 mice with B16F10-OVA cells subcutaneously. Within a week subcutaneous melanomas were palpable. At that time point, we intravenously injected mice with a single cycle LCMV-vector carrying the ovalbumin antigen. In the absence of growing tumors, immunization with LCMV-OVA led to fast expansion of OVA-specific CD8^+^ T cells (Fig. 1A). Interestingly, presence of growing tumors accelerated the expansion of OVA-specific CD8^+^ T cells (Fig. 1A), while treatment without LCMV-OVA did not lead to measurable OVA-specific CD8^+^ T cell responses. Next, we wondered whether the increased number of OVA-specific CD8^+^ T cells could impact on the tumor growth. To this end, we compared growth of B16F10-OVA tumors in mice combined with and without LCMV-OVA application. Surprisingly LCMV-OVA treatment did not affect tumor growth (Fig. 1B).

**Figure 1:**
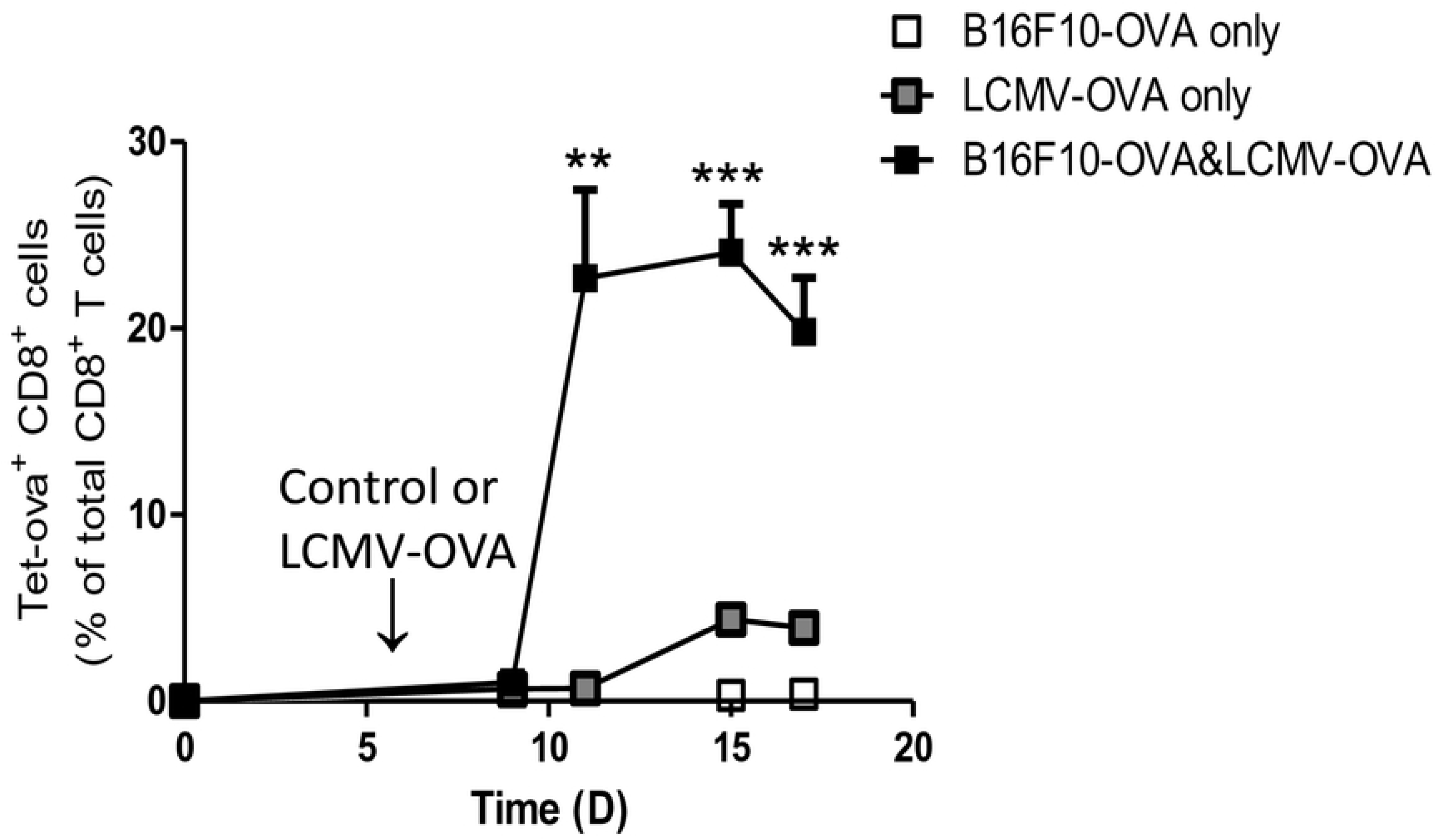
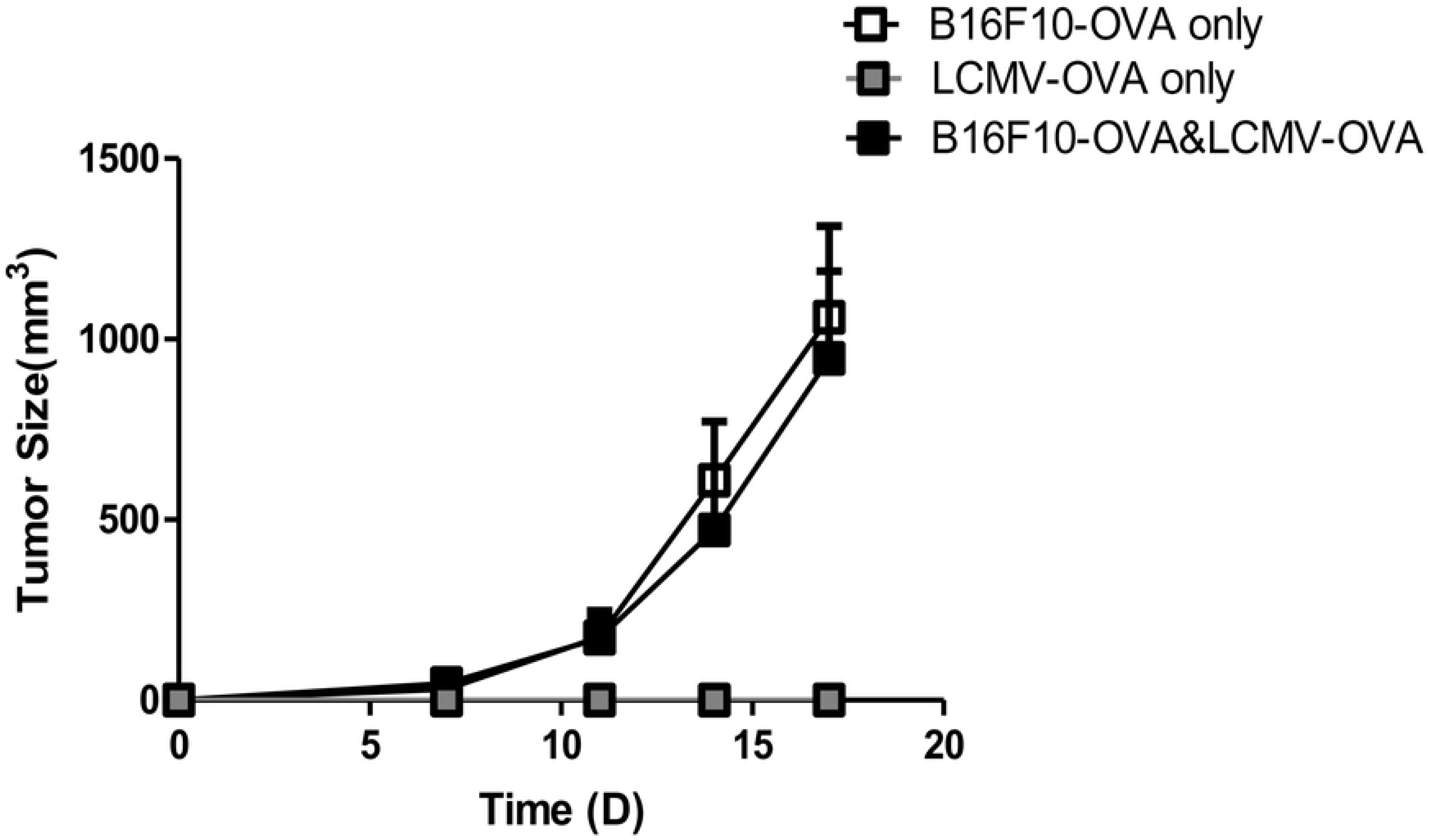
Systemic expansion of tumor-specific CD8+ T cells despite growing tumors. C57BL/6 mice were treated with 1×10^6^ B16F10-OVA cells subcutaneously on day 0. Mice were immunized with 2×10^5^ PFU of LCMV-OVA on day 7. **A:** Tumor growth was analyzed (n = 4). **B:** Frequencies of Tet-OVA^+^ CD8^+^ T cells were analyzed in the blood (n = 4).

### Presence of growing tumors did not influence CD8+ T cell function

Next we aimed to analyze whether the localization and/or function of OVA-specific CD8^+^ T cells could explain unaffected tumor growth in LCMV-OVA treated mice. Therefore, we performed tetramer staining of T cells isolated from spleen, tumor-draining lymph nodes and tumors 10 days after LCMV-OVA immunization. Frequencies of Tet-OVA^+^ CD8^+^ T cells were significantly increased in spleens and lymph nodes of mice, which had received subcutaneous B16-OVA melanomas in addition to LCMV-OVA (Fig. 2A). In line IFN-gamma production of T cells from secondary lymphoid organs after in vitro restimulation was enhanced in B16-OVA bearing mice, which were immunized with LCMV-OVA (Fig. 2B). These data further supported that presence of B16-OVA tumors rather enhanced the anti-tumor immune response than suppressing it. From these data it remains unclear, why tumors still grow in the presence of tumor-specific CD8^+^ T cells.

**Figure 2:**
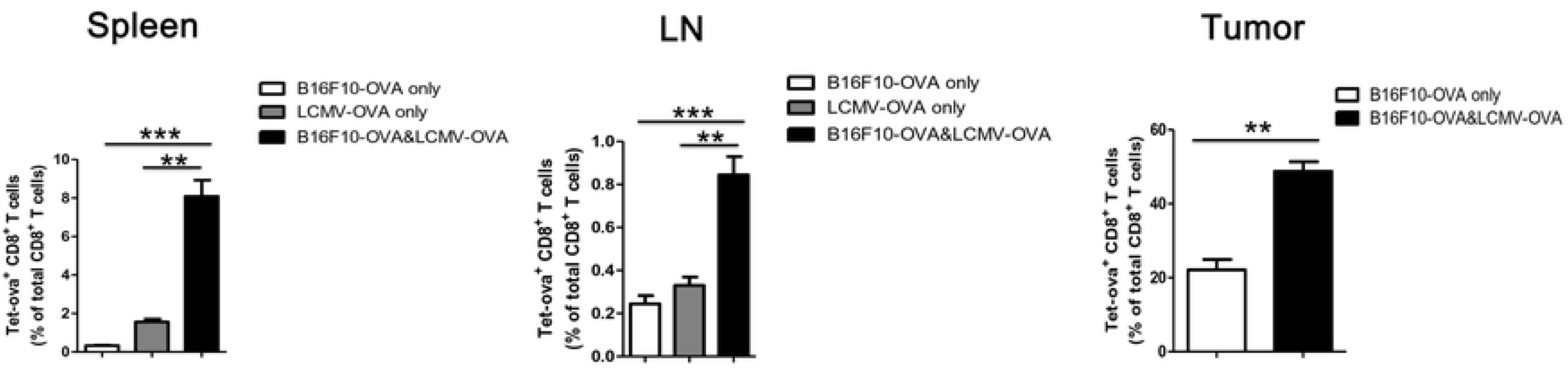
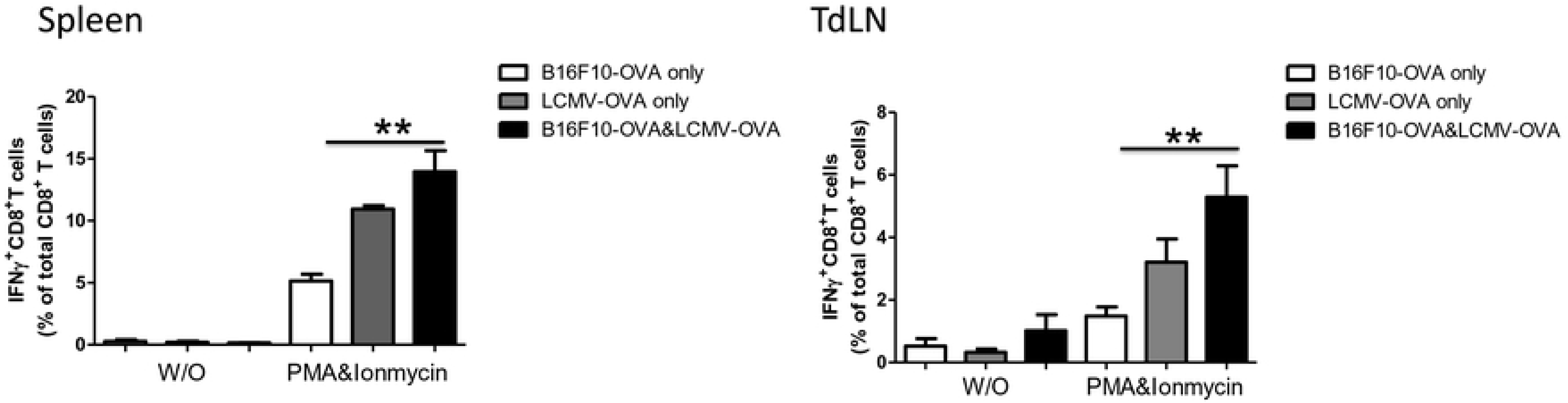
Presence of growing tumors do not influence CD8+ T cell function. C57BL/6 mice were treated with 1×10^6^ B16F10-OVA cells subcutaneously on day 0. Mice were immunized with 2×10^5^ PFU of LCMV-OVA on day 7. **A:** Frequencies of Tet-OVA^+^ CD8^+^ T cells were analyzed in spleen, tumor draining lymph Node and tumor on day 17 (n = 4). **B:** Frequencies of IFN-gamma producing CD8^+^ T cells were analyzed in spleen and tumor draining lymph Node after in vitro restimulation on day 17 (n = 4).

### Tumor-specific CD8+ T cells show normal cytotoxic function in vivo

Next, we wondered whether the systemic and local cytotoxicity is influenced by growing tumors. Therefore we infected mice with LCMV-OVA and then on day 7 injected B16-OVA cells subcutaneously. 24 hours later the amount of injected tumor cells within the injection area were quantified. LCMV-OVA pretreated mice reduced the number of subcutaneously detectable tumor cells (Fig. 3A). This confirmed the anti-tumoral capacity of OVA-specific CD8^+^ T in immunized mice. Next we analyzed whether the presence of B16-OVA tumor cells reduces the cytotoxicity of OVA-specific CD8^+^ T cells. Therefore we treated mice with LCMV-OVA. One group of mice received additionally B16-OVA cells. Cytotoxicity was analyzed after 24 hours by injecting CFSE labeled peptide loaded splenocytes. Presence of B16-OVA cells did not influence specific lysis of OVA-loaded splenocytes within the draining lymph node (Fig. 3B). However, presence of B16-OVA cells accelerated cytotoxicity of OVA-specific CD8^+^ T cells within the injection site (Fig. 3B). Next we asked whether bigger tumors similarly accelerate local CD8 T cell responses. To do so, C57BL/6 mice were treated with B16F10-OVA cells subcutaneously on day 0. Mice were immunized with 2×10^5^ PFU of LCMV-OVA on day 7 and in vivo cytotoxicity was determined eight days later. Circulating OVA-specific T cells from mice harboring growing subcutaneus melanomas showed enhanced cytotoxicity in blood when compared to control animals (Fig. 3C). Together these data suggest, that in our model, growing tumors rather enhanced the frequencies and cytotoxic activity of tumor-specific CD8^+^ T cells than limiting it.

**Figure 3:**
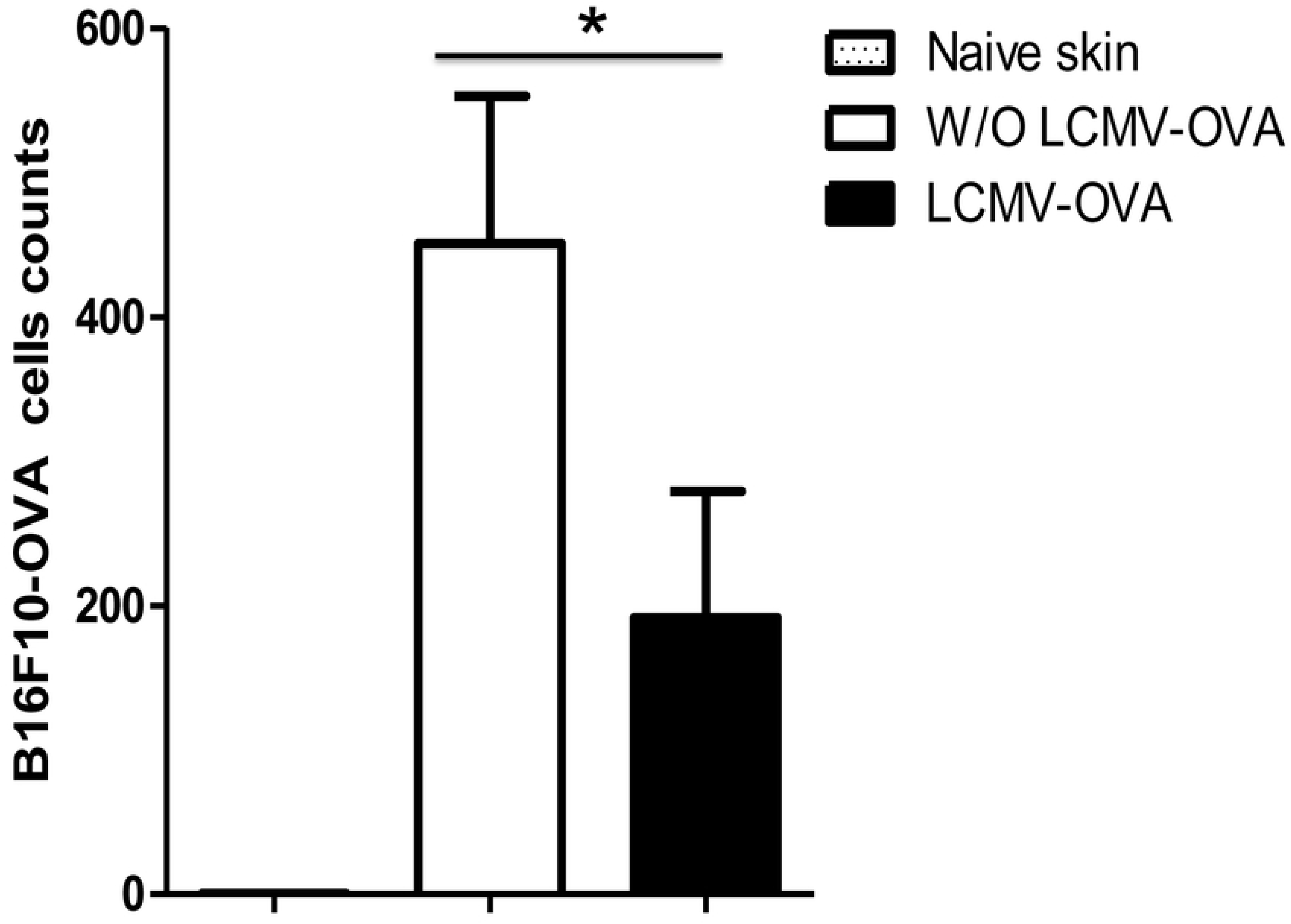
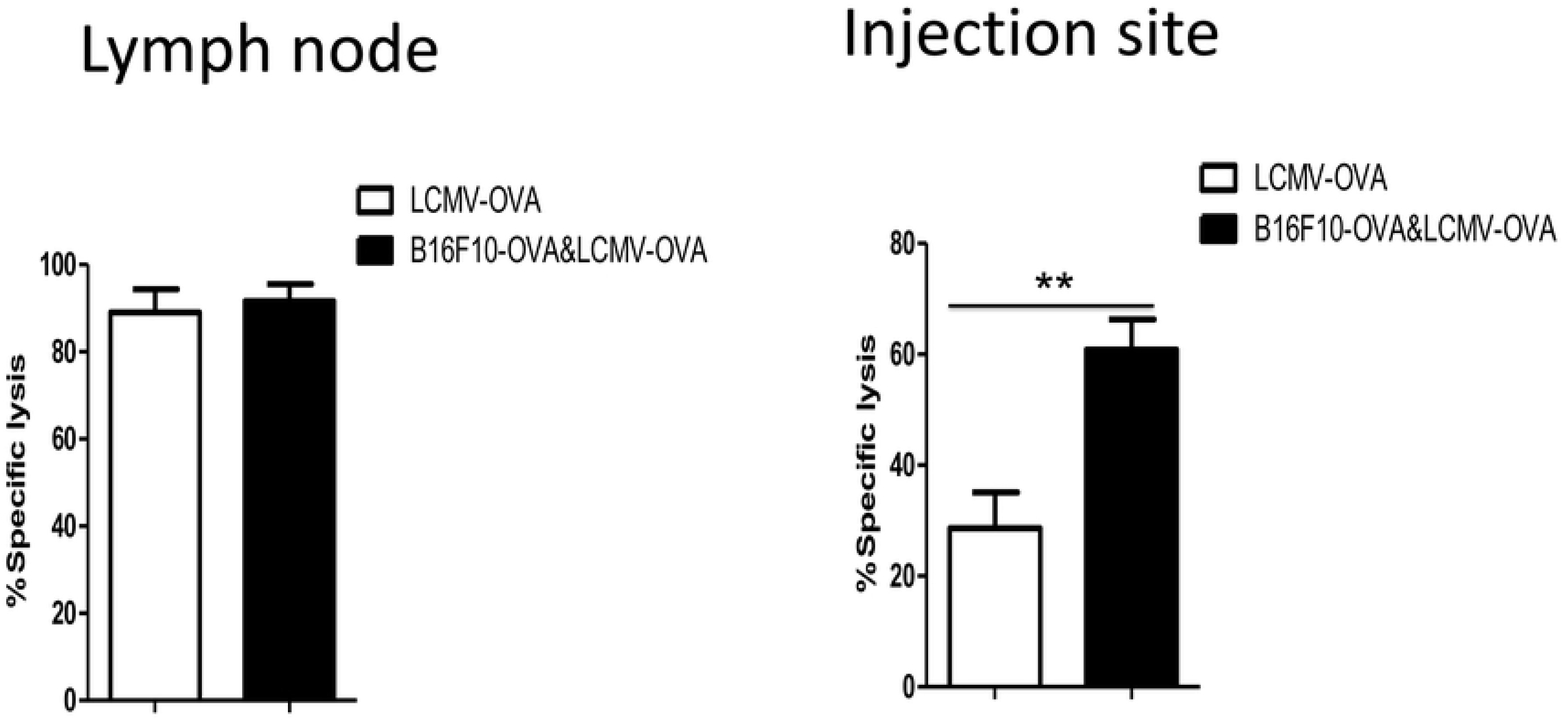
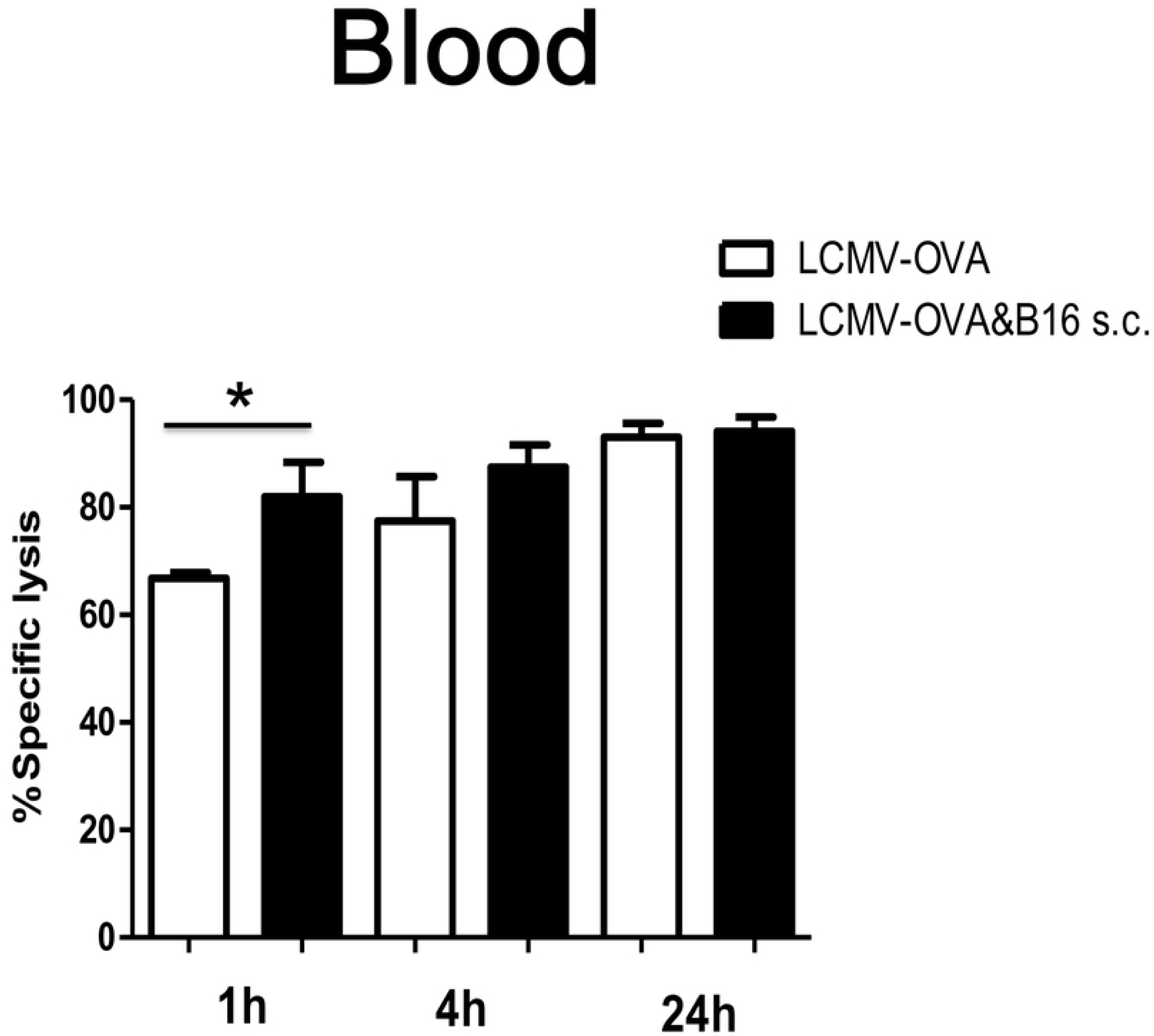
Local tumor-specific CD8+ T cells are cytotoxic. **A:** C57BL/6 mice were treated without (Control) or with 2×105 PFU LCMV-OVA on day 0. On day 7 mice were treated with 1×106 B16F10-OVA cells subcutaneously. Number of B16F10-OVA cells in the subcutaneous injection area was analyzed on day 8 (n = 4). **B:** C57BL/6 mice were treated without (Control) or with 2×10^5^ PFU LCMV-OVA on day 0. On day 7 B16F10-OVA cells were injected subcutaneously. In vivo cytotoxicity was analyzed 24 hours later (n=4). **C:** C57BL/6 mice were treated with 1×10^6^ B16F10-OVA cells subcutaneously on day 0. Mice were immunized with 2×10^5^ PFU of LCMV-OVA on day 7. In vivo cytotoxicity was analyzed on day 15 in blood and the tumor (n=4).

### Growing melanomas down-modulate MHC-I in vivo

So far, our data suggests increased anti-tumoral activity of systemic CD8^+^ T cells in LCMV-OVA immunized mice. However, established subcutaneous tumors progressed despite the activation of cytotoxic immune defenses. We wondered, if the intratumoral antigen-presentation was altered and thus responsible for the lacking effects of cytotoxic T cells on the progression of tumors. To get insights, we performed immunofluorescence of tumor cells or tumors from B16F10-tumors bearing C57BL/6 mice. Indeed, advancing tumors showed limited MHC-I expression compared to tumor cells in culture (Fig. 4).

**Figure 4:**
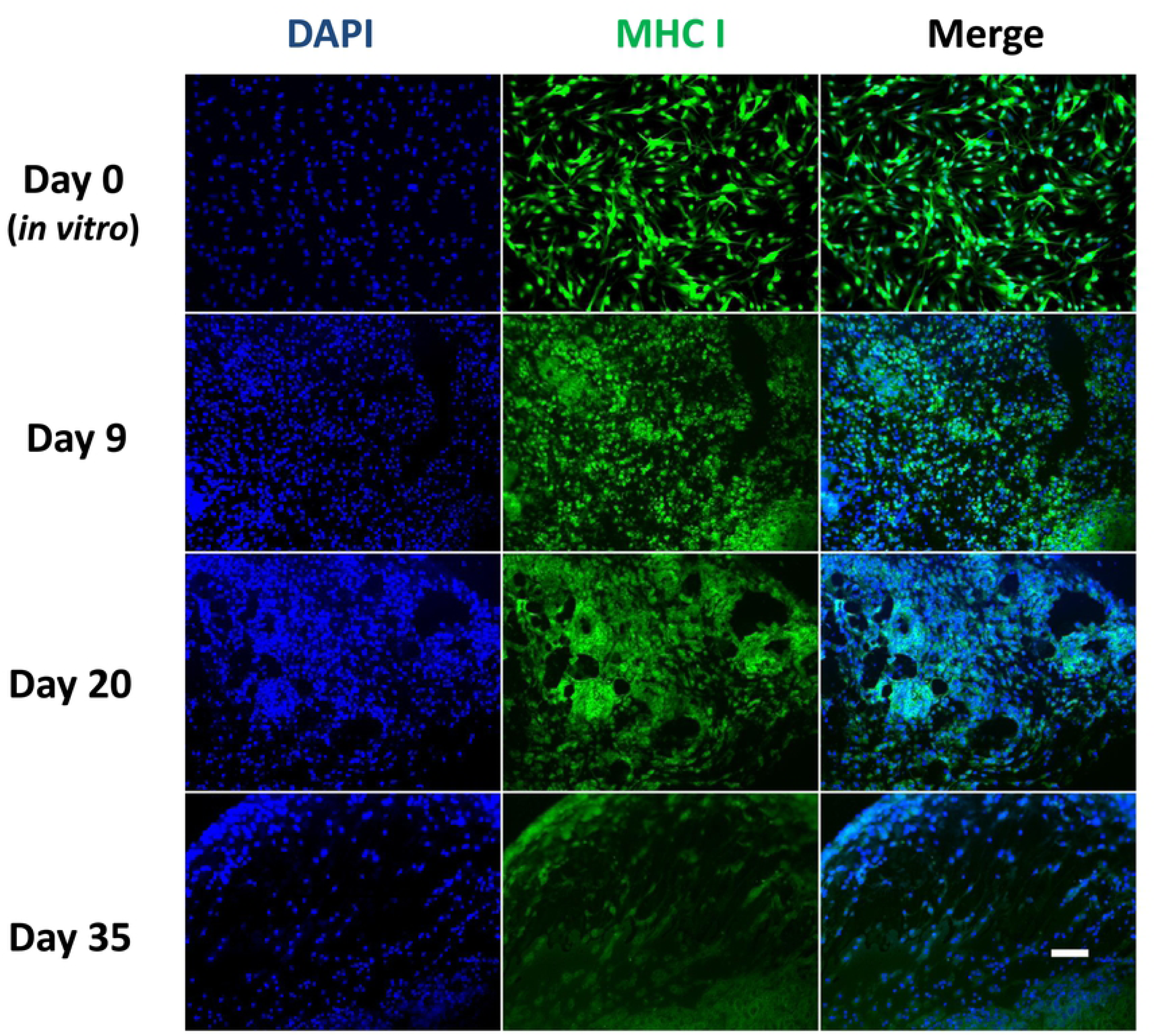
Growing melanoma cells down-modulate MHC-I. Immunofluorescence of tumor cells or tumors from B16F10-tumors bearing C57BL/6 mice (n = 4). Scale bar, 200μm.

### Innate activation by Candid#1 restored MHC-I expression, activated tumor specific CD8+ T cells and limited tumor growth

We postulated that lack of inflammatory signals within the tumor led to limited MHC-I expression. Indeed, infection of tumors with the replicating arenavirus strains LCMV or Candid#1 increased expression of MHC-I in B16 tumors (Fig. 5A). In line with these findings naïve tumor-specific CD8^+^ T cells proliferated and expanded upon LCMV infection (Fig.5B+C). Moreover, virus infection led to an increase of tumor-infiltrating CD8^+^ T cells (Fig. 5D). Since, replication-proficient viruses induce a strong innate immune response that cooperates with adaptive immune defenses, we reasoned, that additional infection with a proliferative, non-cytoloytic arenaviruses might reverse the T cell anergy that we observed in our vaccination model with the single-cycle LCMV-OVA virus. To this end, we again treated C57BL/6 mice with B16F10-OVA cells. One group of mice was treated with LCMV-OVA. Another group was treated with LCMV-OVA and additionally treated with Candid#1. Indeed, Candid #1 treated mice led to significant delay in tumor progression (Fig. 5E).

**Figure 5:**
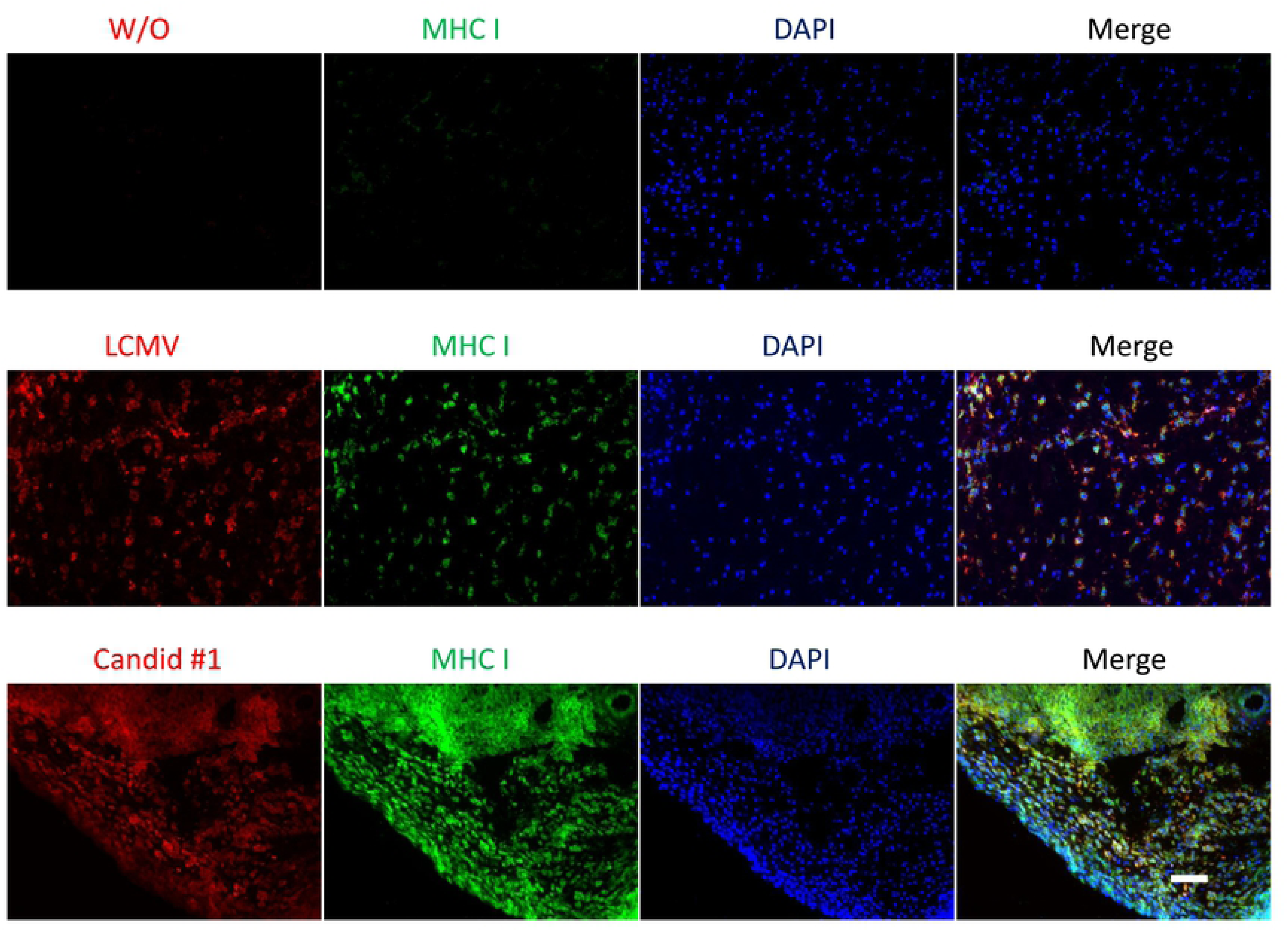
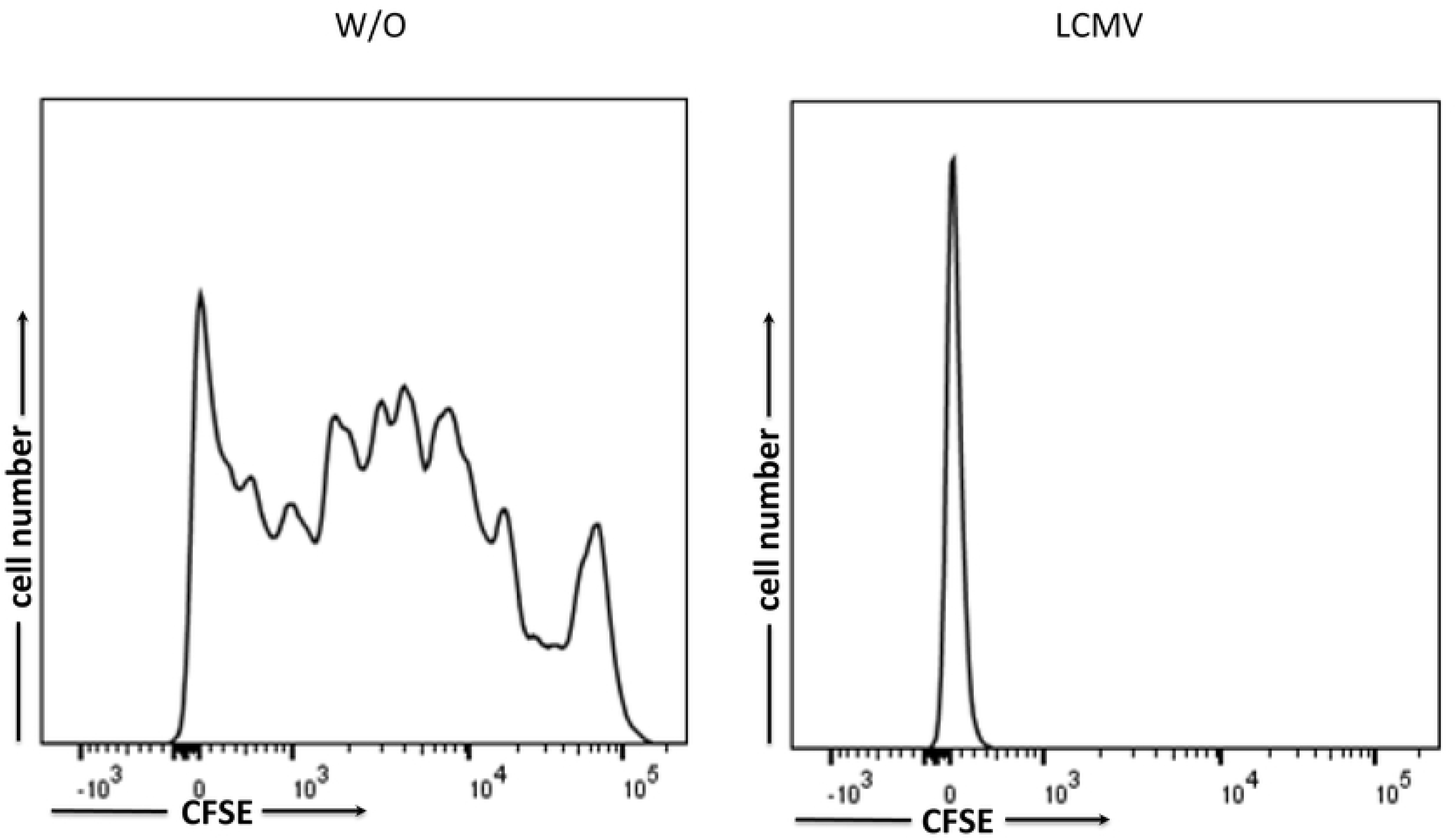
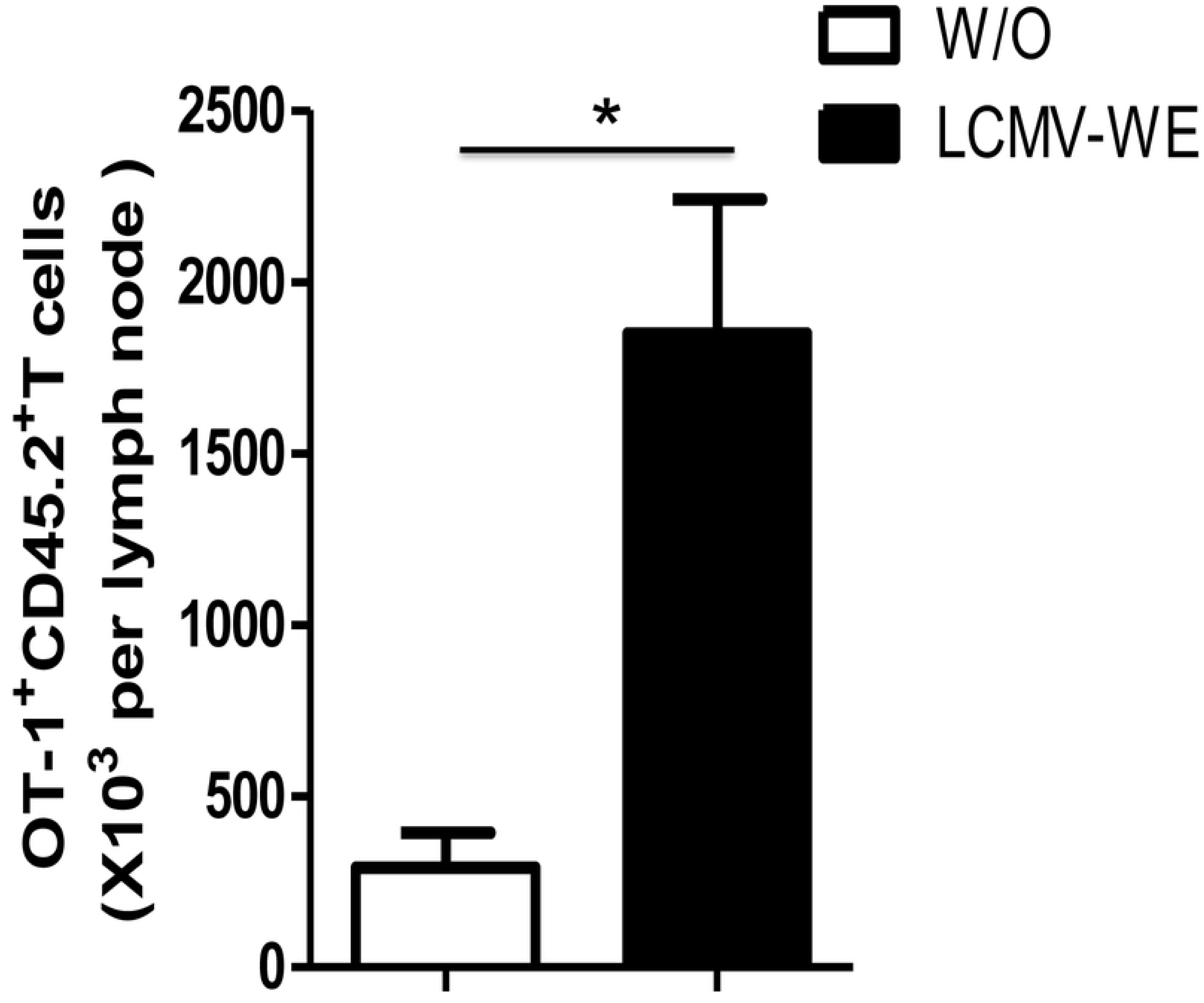
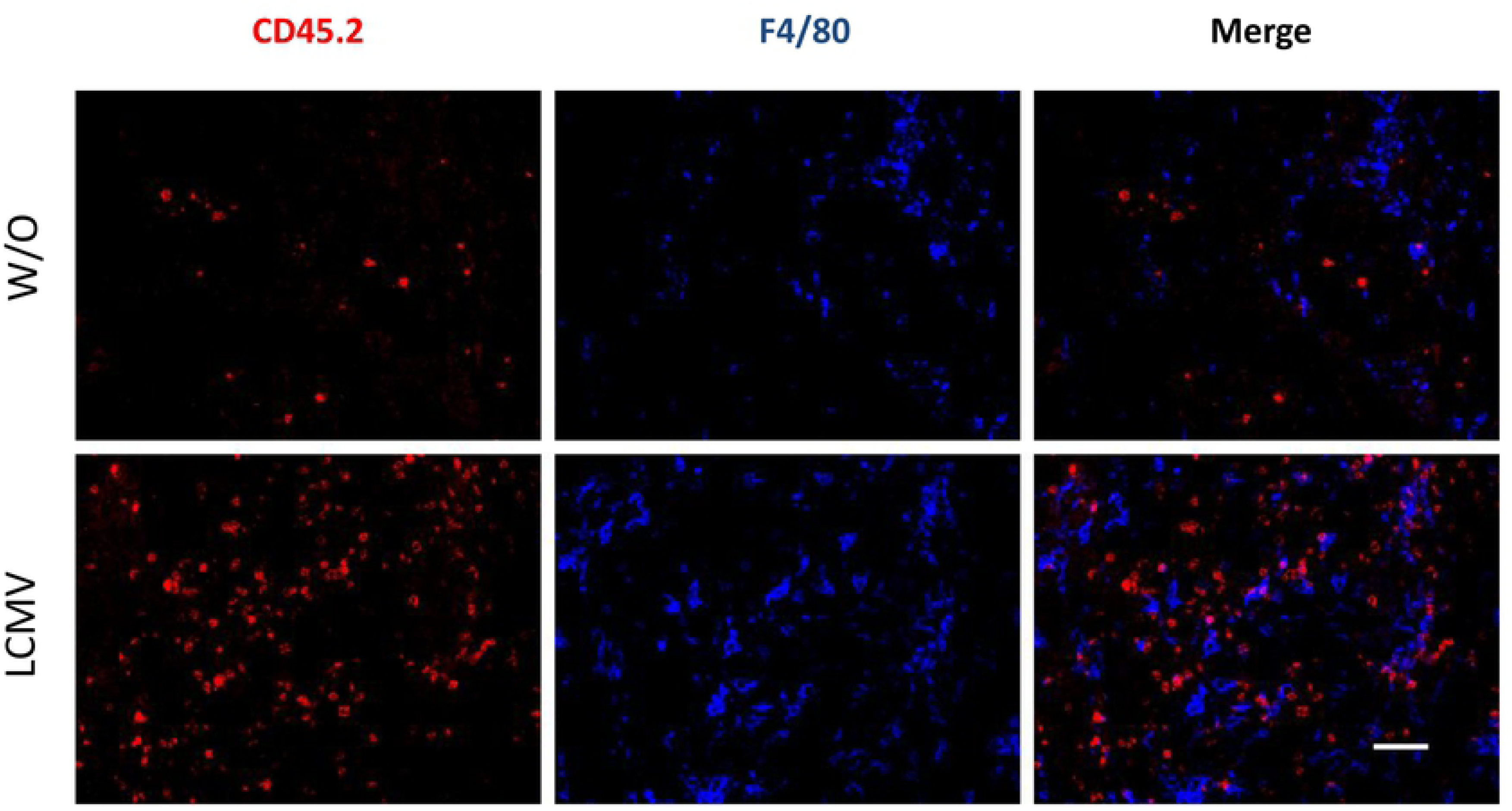
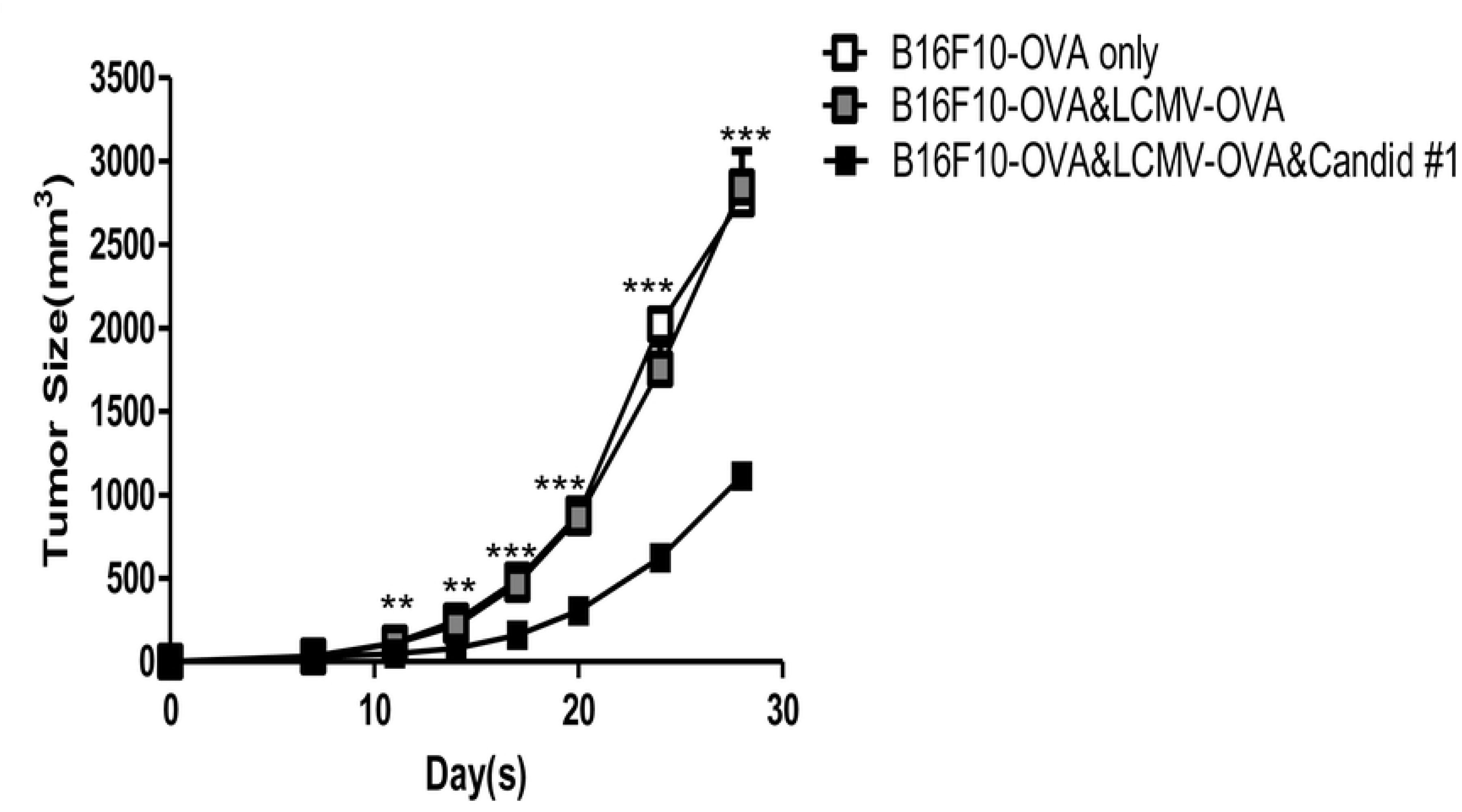
Innate activation by Candid#1 restored MHC-I expression, activated tumor specific CD8+ T cells and limited tumor growth. **A:** C57BL/6 mice were treated with B16F10-OVA cells on day 0. Mice were treated with LCMV or Candid#1 and day 7 and MHC-I expression was determined on day 20 (n = 4). Scale bar, 200μm. **B-D:** CD45.1 mice were treated with B16F10-OVA cells on day 0. On day 7 CFSE-labeled OT-1 cells (CD45.2) were injected intravenously and mice were infected with LCMV on day 8. Proliferation of OT-1 cells in the draining LNs was analyzed on day 13 by CFSE dilution later (**B**, n = 4). Total number of tumor-specific CD8^+^ T cells was determined in the draining lymph node (**C**, n = 4). Infiltration of tumor-specific CD8^+^ T cells was determined by Immunofluorescence (**D**, n = 4). **E:** C57BL/6 mice were treated with B16F10-OVA cells on day 0. Mice were treated with LCMV or Candid#1 and day 7 and tumor growth was monitored.

## Discussion

Our studies focused on the functionality of cytotoxic T cells in a murine melanoma model with B16F10-OVA. We found that vaccination induced expansion and activation of cytotoxic T cells was strongly enhanced when mice where harboring OVA-antigen expressing cancers. However, adaptive immune activation could not prevent or delay progression of established tumors, despite tumor-infiltration. Strikingly, ex vivo restimulation of OVA-specific cytotoxic T cells proved their functionality. Yet, intratumoral cytotoxic T cells remained unresponsive at late stages of tumor. We show that this T cell anergy is associated with low expression level of MHC class I on tumor cells. Furthermore, replicating non-oncolytic arenaviruses like LCMV and Junín virus vaccine, Candid#1 reverse the expression of MHC class I and lead to suppressed tumor growth.

Several types of tumors are characterized by a strong infiltration of cytotoxic T cells reflecting a pre-existing host immunity directed against cancer cells[38]. However, under the circumstances of chronic inflammation, the tumor infiltrating cytotoxic T-cells lose their functionality and undergo exhaustion[4, 39, 40]. Many immunosuppressive factors secreted by tumor cells as well as by immune cells from the tumor microenvironment, such as IL-10, iNOS, ROS or TGF-β are implicated in the cytotoxic T cell dysfunction[41, 42]. PD1-receptor inhibits the reactivity of the cytotoxic T cells converting them to an exhausted phenotype[43, 44]. Consequently, drugs blocking the PD-1 or the PD-L1 restore the anti-tumor activity of the cytotoxic T cells resulting in reduction of tumor growth and clinical benefit in a vast array of tumors of different origin[38, 45-47]. Especially, in the late stage of cancer development PD-1 expression the tumor cells increase causing tumor progression and metastasis due to PD-1 mediated cytotoxic T cell exhaustion[38]. Accordingly, we also saw tumor-specific CD8^+^ T cells infiltrating into advanced melanoma but being anergic to the cancer cells. We assume, that the unresponsiveness of the CD8^+^ T cells to the tumor might be mainly attributed to the downregulation of the MHC class I molecules on the surface of tumor cells. In the late phase, established tumors display less MHC-I on their surface due to the alterations of the components of antigen presentation pathway over time making the malignant cells invisible for the cytotoxic T cells and resulting in immune escape[4, 23]. Nevertheless, reduction of MHC-I expression is a reversible state. Type I and type II interferon signaling contribute to the upregulation of MHC-I on tumor cells[28, 48-50]. In our previous studies[9] we showed that replicating arenaviruses induce a strong interferon response via the activation of the innate immune response. Therefore, we reasoned that arenavirus infection can reverse T cell anergy towards tumor cells via modulation of antigen-presentation. Indeed, our data show an upregulation of MHC I upon viral infection that is accompanied with increased T cell activation in tumors, resulting in suppression of tumor growth.

However, the modulation of the tumor microenvironment and immune-responses within caused by replicating non-oncolytic viruses are very complex. Our study focuses on CD8^+^ T cell activation and observations on MHCI presentation in virus-treated or -untreated tumors. Additional factors to consider, are wide-ranging effects of Interferon signaling induced by viral replication and by activated T cells. These include direct cytotoxic effects of interferons, their impact on tumor-angiogenesis and immunomodulatory effects (REF). Other immune cell types e.g. like dendritic cells, regulatory T cells CD4^+^ T cells might probably contribute to viral infection induced immunomodulation to reverse anergy of cytotoxic T cell and remain subject for future studies.

B16F10-OVA cells activated cytotoxic CD8^+^ T cells. The low amounts of CD8^+^ T cells infiltrating in tumor and the low expression level of MHC class I on tumor surface might explain the failure of cytotoxic T cells regulating the tumor growth. Using an experimental model of melanoma, our results revealed that accumulation and activation of intratumoral CD8^+^ T effector cells was enhanced by LCMV infection, suggesting that the limited infiltration of CD8^+^ T cells observed in B16F10-OVA melanoma recovered by LCMV. We have previously shown that LCMV can replicate in many tumor models[9], which strongly support that LCMV and Junín virus vaccine, Candid#1 reversed the MHC class I expression. We discovered that LCMV treatment promoted the proliferation of CD8^+^ T cells in draining LNs, and LCMV combined PD-1 cocktail significantly reduced tumor growth compared to LCMV treatment alone (data not show). To better understand antitumor immunity, it is very important to explore how LCMV, might be other non-oncolytic arenavirus in the future, affect the function and proliferation of CD8^+^ T cell. Therefore, it would be of great interest to examine the ability of LCMV to induced CD8^+^ T cells function.

In summary, replicating arenaviruses LCMV and Junín virus vaccine, Candid#1 can reserve MHC class I expression in established tumors to overcome T cell anergy. This virus-induced functional immunomodulation might serve as a potential therapeutic strategy for cancer treatment.

